# Simultaneous adjunctive treatment of malaria and its co-evolved genetic disorder sickle cell anaemia

**DOI:** 10.1101/2022.09.01.506230

**Authors:** Innocent Safeukui, Russell E. Ware, Narla Mohandas, Kasturi Haldar

## Abstract

Effective treatments for genetic disorders that co-evolved with pathogens require simultaneous betterment of both conditions. Hydroxyurea (HU) offers safe and efficacious treatment for sickle cell anemia (SCA) by reducing clinical complications, transfusions, and death. Despite concerns that HU-treatment for SCA would increase infection risk by the human malaria *Plasmodium falciparum*, (the genetic driver of the sickle mutation), HU instead reduced clinical malaria. We show that at physiologically relevant exposures, HU (and other ribonucleotide reductase inhibitors) have significant, intrinsic killing activity *in vitro* against blood stages of *P. falciparum*, with low risk of eliciting stably resistant parasites or compromising potency of current antimalarial drugs. Additive activity devoid of antagonism by HU was observed with a wide spectrum of commonly used antimalarial treatments. These data endorse broad, safe, long-term use of HU for SCA in malaria endemic countries and provide a novel biological model for simultaneous, adjunct therapy of a life-threatening infection and concomitant management of a co-evolved genetic disorder.

**Significance:** Genetic disorders are increasingly being treated in global health settings. Hydroxyurea (HU; a ribonucleotide reductase inhibitor) is safe and efficacious for treating sickle cell anemia (SCA). Since the sickle mutation co-evolved with the human malaria parasite *Plasmodium falciparum*, HU-treatment may potentially have increased malarial infection in SCA patients. However, HU reduced clinical malaria, but why this occurred was not understood. We discovered that in doses used in patients, HU kills *P. falciparum* and shows strong potential for safe, adjunctive use with current antimalarial drugs. Our findings endorse, long-term use of HU for SCA in malaria-endemic countries and a novel model for simultaneous, adjunct treatment of a life-threatening infection like malaria with concomitant management of multiple genetic hematological disorders of global proportions.

## Introduction

Treatment of infectious diseases is a major priority in global health^1^. However, inherited genetic diseases also cause significant burden on health care^2^ and therapies to treat them are receiving renewed attention^3^. For the management of genetic disorders that co-evolved with pathogens, it is imperative that new treatments also reduce associated infections, but criteria needed for their evaluation are not well established. Haemoglobinopathies are genetic disorders that are well recognized to be of global importance^4^. Sickle cell anemia (SCA) is an autosomal recessive hemoglobinopathy and one of the most commonly inherited hematological disorders worldwide. SCA is a major global health concern, particularly in sub-Saharan Africa and India where more than 300,000 affected infants are born annually^5^. Recent epidemiological data suggest that over one-third of affected children in sub-Saharan Africa will die before their 5^th^ birthday^6^, often without an established diagnosis. SCA is caused by a 17G>T substitution in the β-globin (*HBB*) gene, which leads to a G6V missense mutation and abnormal polymerization of the resulting sickle hemoglobin (HbS) under deoxygenation conditions, which in turn causes erythrocyte sickling and life-threatening clinical manifestations of severe hemolytic anemia, vaso-occlusive painful crises, stroke and renal disease^7^.

The historic cure of SCA by stem cell transplantation is neither available nor affordable in countries with low resources. However, decades of controlled clinical research have shown that HU, a potent ribonucleotide reductase inhibitor, is both safe and efficacious for SCA and leads to reduced morbidity and mortality, leading to its increasing use to treat both children and adults with SCA in the developed world as well as Africa and India^5,8 9-12^. Long term administration of HU for SCA patients induces production of fetal hemoglobin (HbF), which inhibits erythrocyte sickling, improves red cell rheology, and increases red cell life span^13^. The regions of sub-Saharan Africa and India with high incidence of SCA are also associated with the highest malaria burden^14^. This co-evolution has occurred because a single copy of the *HBB* mutation leads to sickle trait (HbAS), which confers substantial protection against severe malaria and death due to malaria^15^. However individuals who inherit two copies have SCA (HbSS) show lower levels of parasite infection those with normal hemoglobin16 17 but are potentially more susceptible to infection-induced severe anemia and disease^17-19^.

Two large recent clinical trials in different African countries that administered a single daily oral dose of HU to confirm the clinical benefits and safety for SCA also evaluated the effects of HU treatment on malaria^20 21^. The NOHARM study^21^ reported low but similar levels in *P. falciparum* malaria in both treated and placebo group, but all patients received antimalarial prophylaxis with good adherence. In contrast, the REACH study^20^ with many more cases documented ∼50% reduction in malaria infections among the children while receiving hydroxyurea, relative to their pre-treatment rate. The four countries where the REACH trial was conducted, namely the Democratic Republic of Congo (DRC), Angola, Uganda and Kenya show strong co-association between the geographic distribution of SCA and historic *P. falciparum* malaria (ranging from holoendemic to hyperendemic; Fig. 1a-b). The findings from the NOHARM and REACH studies are particularly important because they dispel concerns that long term administration of HU may worsen malaria in children with SCA^22 23 24^, and also showed benefit by reducing malaria incidence and mortality, which are key targets of the 2000 and 2015 Millennium Development Goals.

**Fig. 1.**
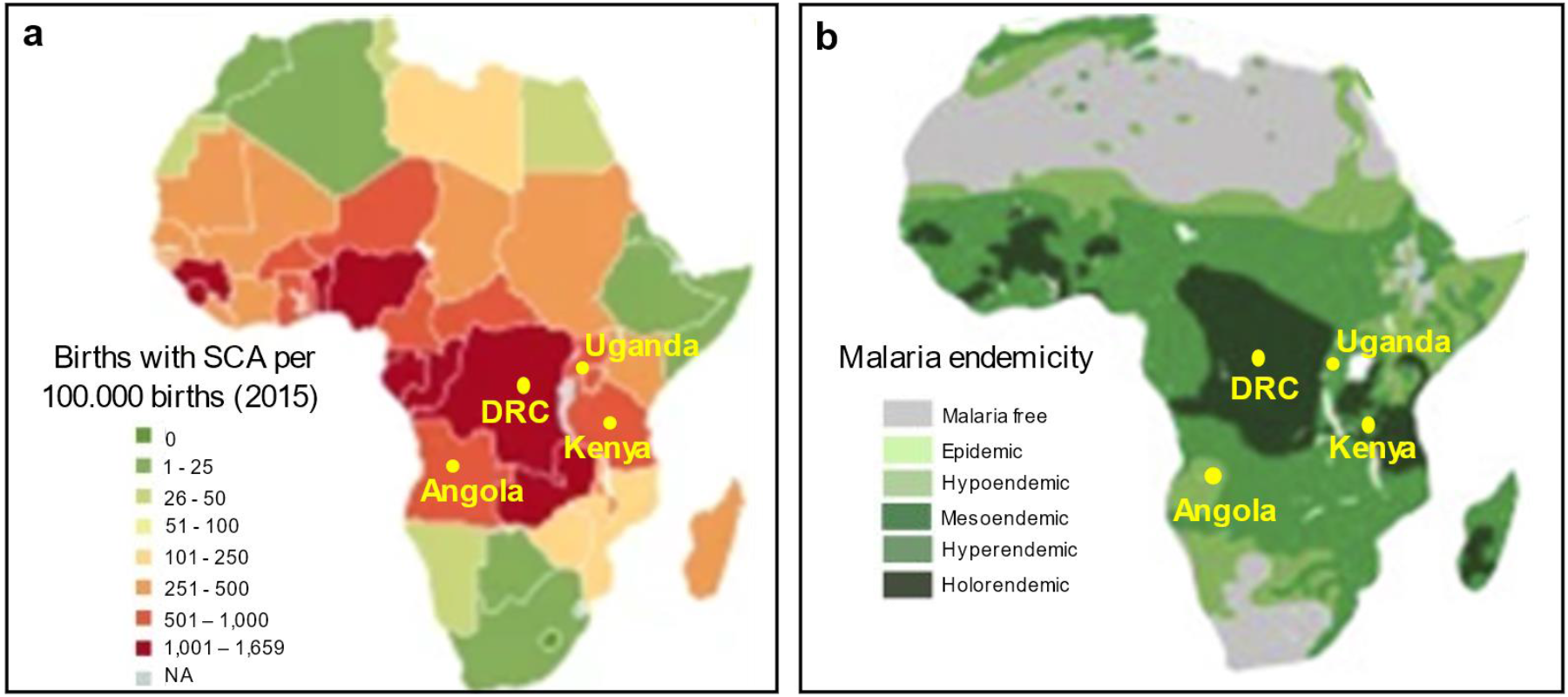
Co-evolution of SCA with *P. falciparum* malaria in Africa. **a-b** Geographical distribution of (a) SCA and (b) *P. falciparum* malaria in Africa (adapted from Kato GJ, Piel FB *et al*., 2018 ^45^ and Piel et al. 2010^14^). Locations of Democratic Republic of Congo (DRC), Uganda, Angola and Kenya countries where the REACH trial are conducted are shown.

We reasoned that the substantial reduction in the incidence of *P. falciparum* malaria may be due to direct anti-plasmodial activity of HU, since orthologues of ribonucleotide reductase (RNR) are encoded by *Plasmodium* species. We therefore explored the intrinsic anti-parasite effects of HU and other RNR inhibitors against laboratory malaria infections. We also queried the effects of HU action on anti-parasite activity of antimalarials used in sub-Saharan Africa, and the efficacy of major antimalarial drugs on HU-tolerant parasites for their long-term implications of potentially decades of SCA treatment. Our findings provide mechanistic understanding by which HU administration directly protects children with SCA against acute and severe malaria, which informs the drug management of SCA and malaria without risk of eliciting parasite drug resistance. They also yield a novel model with potential to be applicable to treatment of an infectious disease like malaria and co-evolved genetic hematological disorders whose control is important for global health.

## Results

### HU and additional RNR inhibitors inhibit *P. falciparum* growth *in vitro*

Since SCA and malaria have co-evolved, drugs used to treat SCA in malaria-endemic regions should be evaluated to ensure they do not promote proliferation of malaria parasites. We investigated the effects of HU exposure on the growth of *P. falciparum* in an *in vitro* culture system. Since HU inhibits RNR we also evaluated three additional known RNR inhibitors: clofarabine (Clof), gemcitabine (Gem) and triapine (3AP) (Fig. 2a). In a standard 72 h treatment, HU inhibited the parasite strain *Pf*3D7, with an EC_50_ of 127.6 µM (Fig. 2b and Table 1). Clof showed a comparable EC_50_ of 100.1 µM but Gem and 3AP were more potent with EC_50_ of 0.8 µM and 0.55 µM, respectively. Similar effects were detected in two additional parasite strains *Pf*NF54 and *Pf*Dd2 (Fig. 2c, Supplementary Fig. 1 and Table 1). The EC_50_ of HU and Gem was higher (1.6 to 1.8-fold) in *Pf*Dd2 (Fig. 2c and Table 1). 3AP was more potent against *P*fDd2 while Clof was equally effective against all three strains. HU’s inhibitory effect was the most modest but it was active across multiple genetic backgrounds. It may be slightly less effective against *Pf*Dd2 strain (which is CQ resistant; but this was not shared across all RNR inhibitors).

**Fig. 2.**
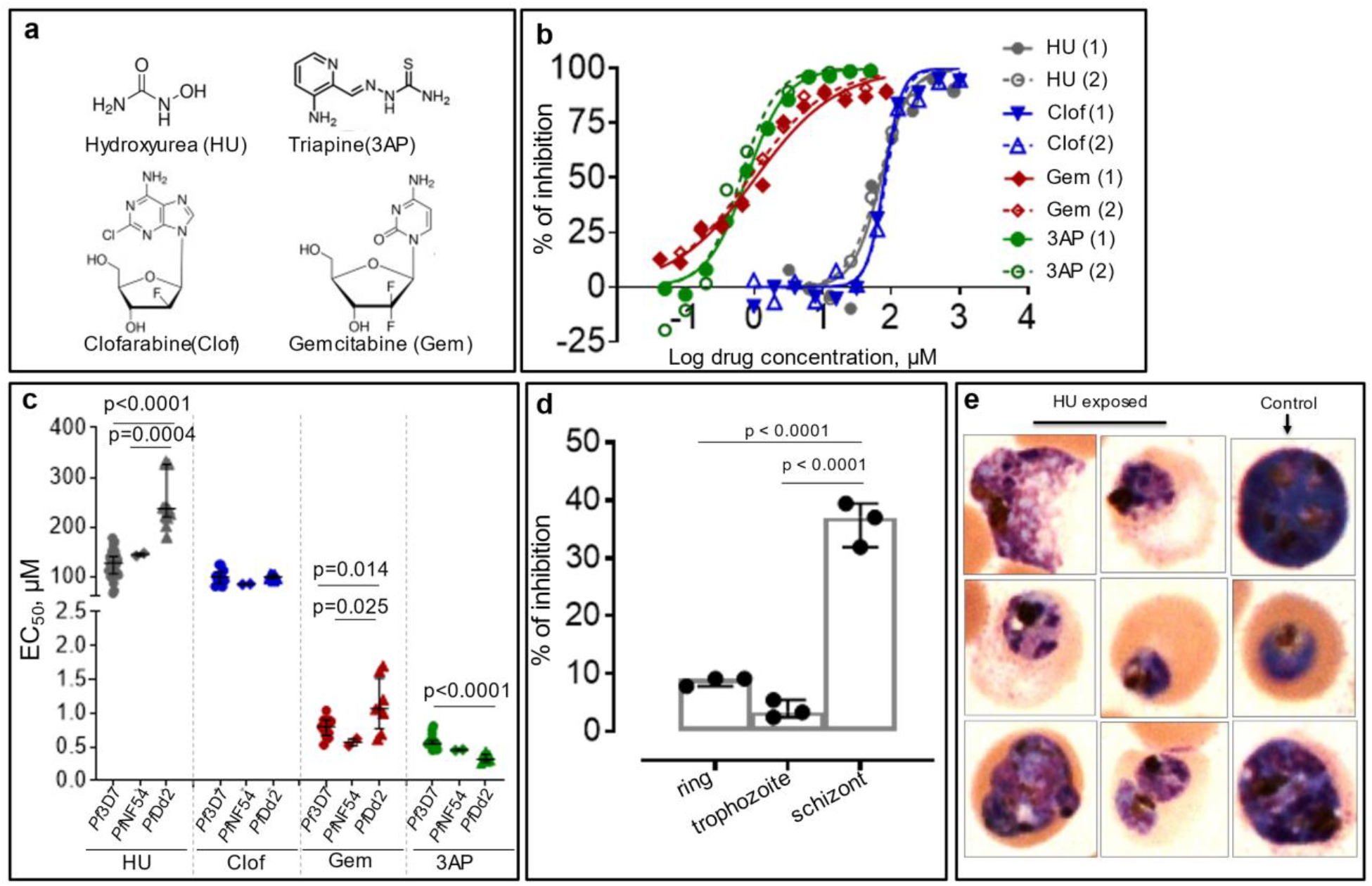
HU and additional RNR inhibitors inhibit *P. falciparum* growth *in vitro* in a stage specific manner. **a** Chemical structure of HU and three additional RNR inhibitors (3AP, clof and gem). **b** Representative inhibitory dose-response curves of *Pf*3D7 strain exposed to indicated drugs for 72 hr (in a standard EC_50_ assay). **c** EC_50_ of HU, 3AP, clof and gem against *Pf*3D7, *Pf*NF54 and *Pf*Dd2 strains. **d** Exposure of *Pf*3D7 ring, trophozoite and schizont stages by a 3h pulse of 329 µM HU and assessment of their development to subsequent stages. **e** Morphological analyses of maturation of rings exposed 72 h to 394.8 µM of HU, as detected by Giemsa staining. In panels c and d, the horizontal bar represents median and vertical bar the interquartile range. For the mean comparisons in panels c and d, one-way ANOVA with a Tukey post hoc analysis was used.

**Table 1.** Effective concentration of HU or other RNR inhibitors, and ACT’s partner drugs or active metabolite of ART against *Pf*3D7, *Pf*NF54 and *Pf*Dd2 asexual blood stages *in vitro*.

| Molecule | *Pf*3D7 |  | *Pf*NF54 |  |  | *Pf*Dd2 |  |  |
| --- | --- | --- | --- | --- | --- | --- | --- | --- |
| | EC<sub>50</sub>, μM | EC<sub>90</sub>, μM | EC<sub>50</sub>, μM | EC<sub>90</sub>, μM | <sup>a</sup>Fold difference | EC<sub>50</sub>, μM | EC<sub>90</sub>, μM | <sup>b</sup>Fold difference |
| **RNR inhibitor** |  |  |  |  |  |  |  |  |
| HU | 127.6 (105.8-140.8) | 471.5 (376.8-592.8) | 144.8 (142.3-147.4) | 344.1 (302.6-459.3) | 1.13 | 225.9 (201.5-235.6) | 759 (499-1043) | 1.77 |
| 3AP | 0.55 (0.54-0.60) | 2.30 (1.7-3.0) | 0.46 (0.45-0.46) | 0.89 (0.81-0.99) | 0.84 | 0.32 (0.28-0.39) | 0.50 (0.46-0.78) | 0.58 |
| Gem | 0.80 (0.67-0.90) | 13.1 (10.4-19.7) | 0.57 (0.52-0.62) | 12.7 (11.1-14.2) | 0.71 | 1.07 (0.77-1.52) | 54.96 (34.64-86.25) | 1.33 |
| Clof | 100.1 (86.8-110) | 189.2 (168-227.2) | 85.2 (84.4-86.0) | 156.4 (148.5-162.1) | 0.85 | 99.8 (96.21-104.08) | 259.4 (227.85-289.4) | 1.00 |
| **Antimalarial drug** |  |  |  |  |  |  |  |  |
| LUM x 10<sup>-3</sup> | 13.1 (5.3-26.8) | 82.1 (31.7-165.4) | ND | ND | - | 15.63 (13.32-17.93) | 57.52 (45.12-69.91) | 1.19 |
| AQ x 10<sup>-3</sup> | 12.9 (9.3-17.4) | 21.6 (15.6-24.1) | ND | ND | - | 14.76 (12.54-16.97) | 32.14 (19.62-44.66) | 1.14 |
| CQ x 10<sup>-3</sup> | 13.6 (11.8-16.7) | 20.3 (18.4-23.4) | ND | ND | - | 100.8 | 193.2 | 7.41 |
| PQ | 0.37 (0.29-0.44) | 2.2 (1.9-2.7) | ND | ND | - | 0.67 (0.55-0.78) | 1.58 (1.20-1.95) | 1.81 |
| TQ | 0.06 (0.04-0.09) | 0.61 (0.32-1.84) | ND | ND | - | ND | ND | - |
| DHA x 10<sup>-3</sup> | 4.1 (3.0-5.2) | 17.2 (9.5-25.3) | ND | ND | - | 4.94 (3.57-6.30) | 11.3 (9.15-13.45) | 1.20 |

Effective concentration (EC): Median (interquartile range) <sup>a</sup>, *Pf*NF54/*Pf*3D7 ratio of EG <sup>b</sup>, *Pf*Dd2/*Pf*3D7 ratio of EG

During blood stage of growth, *P. falciparum* progresses through morphologically distinct, stages, namely rings (formed immediately after parasite entry in the red cell), trophozoites (that develop after 17-24 h), followed by the multinucleated schizont stage. To determine which parasite stages were targeted by HU, each stage was separately exposed to drug. Further, parasites were treated with a bolus of HU to better mimic its daily pharmacological exposure at high micromolar levels in SCA patients. Based on human pharmacokinetic studies ^25 26^, we used 329 µM of HU for 3 hours, after which the drug was washed out and parasites were allowed to mature for additional 69 hours. As shown in Fig. 2d, ring and trophozoite stage parasites were inhibited by <10%. However, growth and proliferation of schizonts was more substantially blocked by 37%. Together, these data confirmed that HU (and additional RNR inhibitors) had intrinsic activity against *P. falciparum* principally by blocking schizonts (Fig. 2e), which is consistent with the requirement of RNR and high levels of DNA synthesis at this stage of parasite development. Notably, peak blood concentrations of HU achieved in children with SCA often exceed 300 µM ^25^, suggesting daily exposure to HU alone may be detrimental to parasites.

### Evaluation of the interaction of HU and additional RNR inhibitors with existing antimalarial drugs

A second key consideration on the potential wider use of HU is its potential interaction with currently used antimalarials. ACTs are the leading antimalarials that reduce parasite burden as well as death due to malaria and are central to malaria elimination worldwide. They are composed of fast-acting artemisinins as well as a slow-acting partner drug. Artemisinins potently inhibit ring as well as later trophozoite and schizont parasite stages. But they are also rapidly cleared *in vivo* (t_1/2_ < 30 min) and must be administered multiple times even during a short 3-day antimalarial treatment course. In contrast, the partner drugs can remain in circulation for months. Long-lived drugs are more likely to have greater interactions with HU administered daily for months to years.

We therefore analyzed interactions of HU with a wide range of antimalarials. We used a standardized, fixed-ratio method used to study drug-drug interactions of combinations^27^. Our drugs of choice included lumefantrine (LUM) and amodiaquine (AQ), the most widely used ACT partner drugs in Africa. CQ, primaquine (PQ) and tafenoquine (TQ) were also evaluated since they are used in treatment of *P. vivax*, a second malaria parasite that is on the rise. More recently PQ has been used in *P. falciparum*-treatment to block gametocytes (the sexual stage in the blood that is transmitted to the mosquito^28^). We also assessed dihydroartemisinin (DHA), the active metabolite of all artemisinin drugs that are used to treat both *P. falciparum* and *P. vivax*.

EC_50_ data for all antimalarial drugs needed to deploy the fixed ratio method are summarized in Table 1. Briefly, the ring stage of *Pf*3D7 and *Pf*Dd2 strains were cultured for 72 h in the presence of different concentrations and proportions of two test combinations (HU and an antimalarial) over the indicated times. We additionally tested the effects of other RNR inhibitors, Gem, Clof and 3AP (in place of HU). The observed values for EC_50_ and EC_90_ were analyzed in relation to the data obtained with a single combination^27^. The fractional inhibitory concentrations (FICs) of HU, other RNR inhibitors and the currently used antimalarial drug at the EC_50_ and EC_90_ were then calculated and plotted in isobolograms. Sums of the FICs (**∑**FICs) of the drug combinations were calculated to evaluate different degrees of inhibition at EC_50_ and EC_90_ and used to infer drug-drug interactions as previously described^29^: ∑FIC < 0.5 indicates substantial synergism, ∑FIC <1 indicates moderate synergism, ∑FIC ≥1 and <2 indicates additive interaction, ∑FIC ≥2 and <4 indicates slight antagonism and ∑FIC > 4 indicates marked antagonism. As shown in Fig. 3a-b and Supplementary Fig. 2, the median **∑**FICs predict additive effects of HU with LUM, AQ, CQ, PQ, TQ and DHA. The range associated with **∑**FICs suggest that there may be trends toward synergy or slight antagonism with LUM, PQ and TQ, but these are expected to be minor (Fig. 3b, see footnote). Based on median **∑**FICs additive effects are also predicted for other RNR inhibitors with these antimalarials (Supplementary Fig. 2). This was observed in the *Pf*3D7 strain as well as the CQ resistant *Pf*Dd2 strain, suggesting conservation across strains.

**Figure 3.**
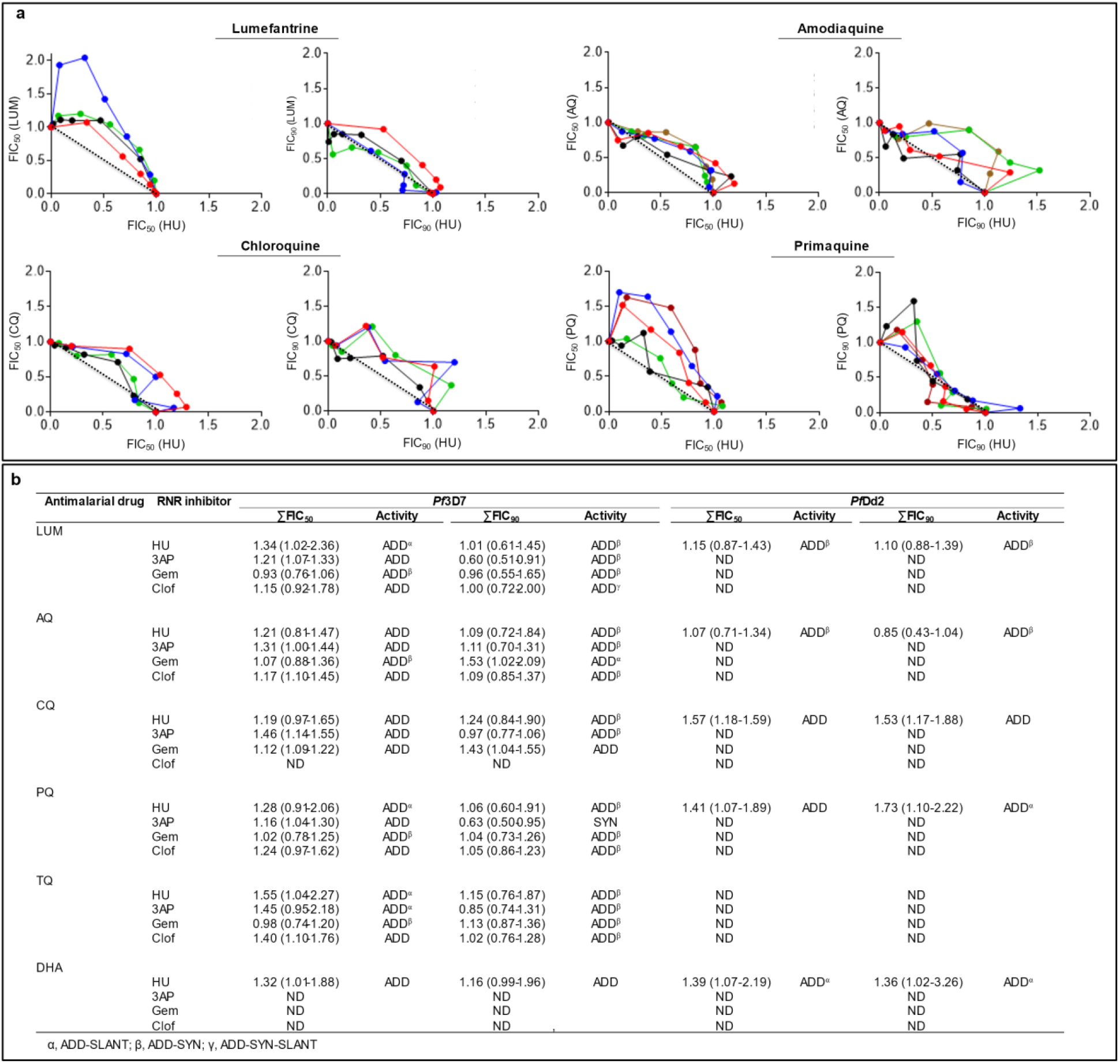
Analyses of the interactions between HU and antimalarial drugs. **a** Isobologram analyses of the interaction between HU and indicated antimalarial drugs against *Pf*3D7: lumefantrine (LUM), amodiaquine (AQ), chloroquine (CQ) and primaquine (PQ). Fractional inhibitory concentrations (FIC) of HU *versus* FIC of antimalarial drugs were plotted. Isobolograms were constructed from EC_50_ or EC_90_ values. For each drug combination, the FIC were calculated by dividing the measured ‘‘apparent’’ EC values for individual drugs in the different combinations of HU and antimalarial drugs EC values obtained when the drugs were used alone. **b** Sum of 50% and 90% fractional inhibitory concentration (∑FIC50, 90) of the interaction of HU and indicated antimalarial drugs against *Pf*3D7 and *Pf*Dd2 strains, carried out in 2 to 5 independent experiments. In panel b, ∑FIC values are presented as median (range). α, ADD-SLANT; β, ADD-SYN; γ, ADD-SYN-SLANT. ADD, additive; SLANT, slightly antagonistic; SYN, synergistic; ND, not determine.

LUM, AQ, CQ and DHA kill parasites at all blood stage parasites. Hence the additive effect of HU with these drugs is likely due to its added effect on schizonts. However, little is understood about killing effects of 8-aminoquinolines: we were particularly interested in PQ, which remains of high value in targeting both *P. falciparum* and *P. vivax* species and stages that are not blocked by other antimalarials. We therefore re-evaluated dose-response curves of PQ + HU starting at early ring stage (Supplementary Fig. 3a) and their stage-dependent action in short ‘pulsed’ treatments with one or both drugs followed by a longer chase with one or both (Supplementary Fig. 3b). Since HU preferentially targets schizonts, we separately treated ring, trophozoite and schizont stages with a pulse of HU, followed by HU alone or PQ alone (Fig. 4a-f). PQ was less effective at the schizont stage (Fig. 4a-f and Supplementary Fig. 3d). In contrast, but as expected, HU showed better activity against schizonts (Fig. 4a-f and Supplementary Fig. 3c). We therefore inferred that HU and PQ act at different stages to induce the highest parasite killing.

**Figure 4.**
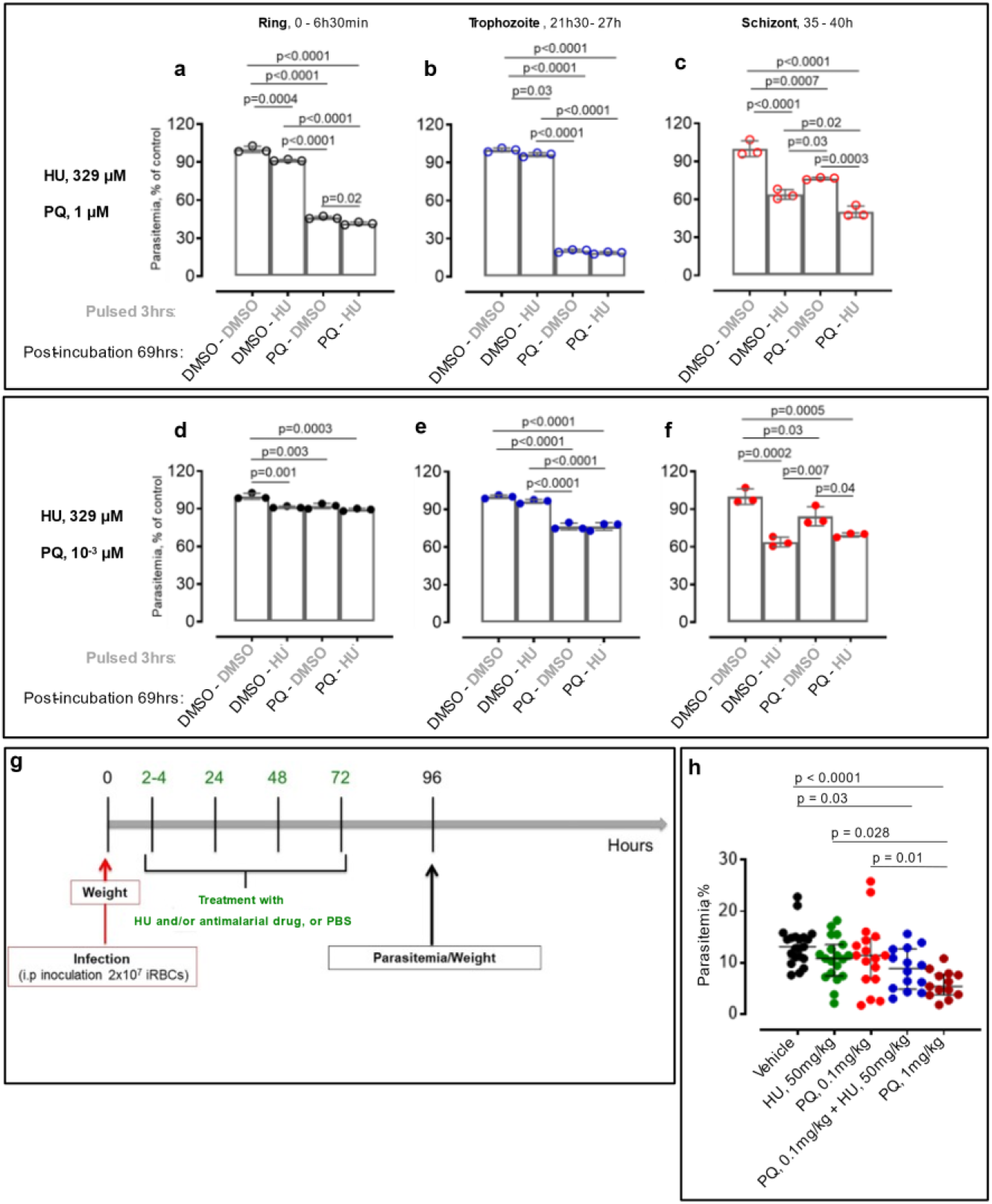
Extended analyses of the anti-plasmodial activity of HU and PQ *in vitro* and *in vivo*. **a-f** Inhibition rate of PQ and/or HU according to the age of asexual blood stage parasites. Parasites (ring, 0 – 6.5h; trophozoite, 21.5 - 27h; schizont, 35 - 40h) were pulsed 3 hours with 329 µM HU or 0.1% DMSO, then extensively washed and incubated 69 hours in media containing 1% DMSO or 329 µM HU or 1 µM (Panels a-c) or 10^−3^ µM (Panels d-f) of PQ. In panels a to f, DMSO-treated parasites were used as controls. **g-h** Additive activity of HU and PQ in *Plasmodium berghei* Anka-infected mice. **g** Schematic representation of *P. berghei* Anka (PbA) infection, drug treatment, body weight and tail blood (for parasitemia) collections. **h** Parasitemia after 4 days of treatment. In panels a-f and h, horizontal bar represents median and vertical bar the interquartile range. For the mean comparisons in panels a-f and h, one-way ANOVA with a Tukey post hoc analysis was used.

To assess whether the observed *in vitro* additive effects of HU and PQ could be observed *in vivo*, we used the *P. berghei* Anka (PbA) infection model in C57BL/6 mice (as summarized in Fig. 4g and Supplementary Fig. 4). Briefly, mice were treated with HU (50 mg/kg) and/or PQ (0.1 or 1 mg/kg), starting 2-4 hours after infection. Drugs were administered every 24 hours for additional 3 consecutive days and parasitemias determined at the indicated time points. As shown in Fig 4g, a combination of HU and PQ (0.1 mg/kg) reduced parasite growth relative to vehicle treatment. But neither drug alone had significant effect although raising the dose of PQ (to 1 mg/kg) was suppressive. In contrast to PQ, the action of TQ in single or multiple doses was not affected by HU (Supplementary Fig. 4 a-b) presumably due to the greater plasma exposure of TQ. At the doses used, HU did not induce weight loss in mice (which was used as the first indication of overt toxicity; Supplementary Fig. 4c-d).

Together these data suggest that HU can contribute additive anti-parasite effects *in vivo* that may help dose management, particularly for antimalarials with lower plasma exposure. The additive effect of HU is likely due to its efficacy in blocking schizonts. Further HU is additive across multiple broad classes of antimalarials with different chemical scaffolds as well as different *P. falciparum* strains.

### Selection and characterization of HU resistant *P. falciparum*

To identify targets of HU in *P. falciparum*, we selected for parasites resistant to the drug. We performed single-step *in vitro* resistance selections by exposing *Pf*3D7 and *Pf*Dd2 parasites to HU in one step of selections at 5× or 3x EC_50_ (Materials and Methods). Both failed to yield resistant parasites after 60 days (data not shown) suggesting neither strain was likely to readily become resistant to HU. However, slow increase of HU over 90 days in four gradual, sequential steps of drug concentrations at 1, 1.5, 3.1 and 6.7 x EC_50,_ led to selection of *Pf*3D7 parasites with EC_50_ greater than 850 µM (3D7HU65, as compared to the starting parasites with EC_50_ of 131 µM; Fig. 5a-c). Under the same chemo-selection scheme over 90 days, *Pf*Dd2 showed a shift of < 2 x EC_50._ (Supplementary Fig. 5). Since *Pf*Dd2 parasites showed such a low propensity to develop resistance, subsequent experiments were limited to the *Pf*3D7 strain. Several independent clones derived from 3D7HU65 in presence of HU, showed EC_50_’s comparable to the bulk population (Fig. 5d-f). Two clones were expanded in drug free media: one was with the highest EC_50_ and the second was with the lowest EC_50._ Both were then exposed to 850 µM HU. Both failed to proliferate (Fig. 5g), suggesting gain of resistance in presence of HU was likely transient (due to plasmid-based amplification of putative resistance genes) rather than stable resistance due mutation or duplication of the parasite’s chromosomal copy of the target gene.

**Fig. 5.**
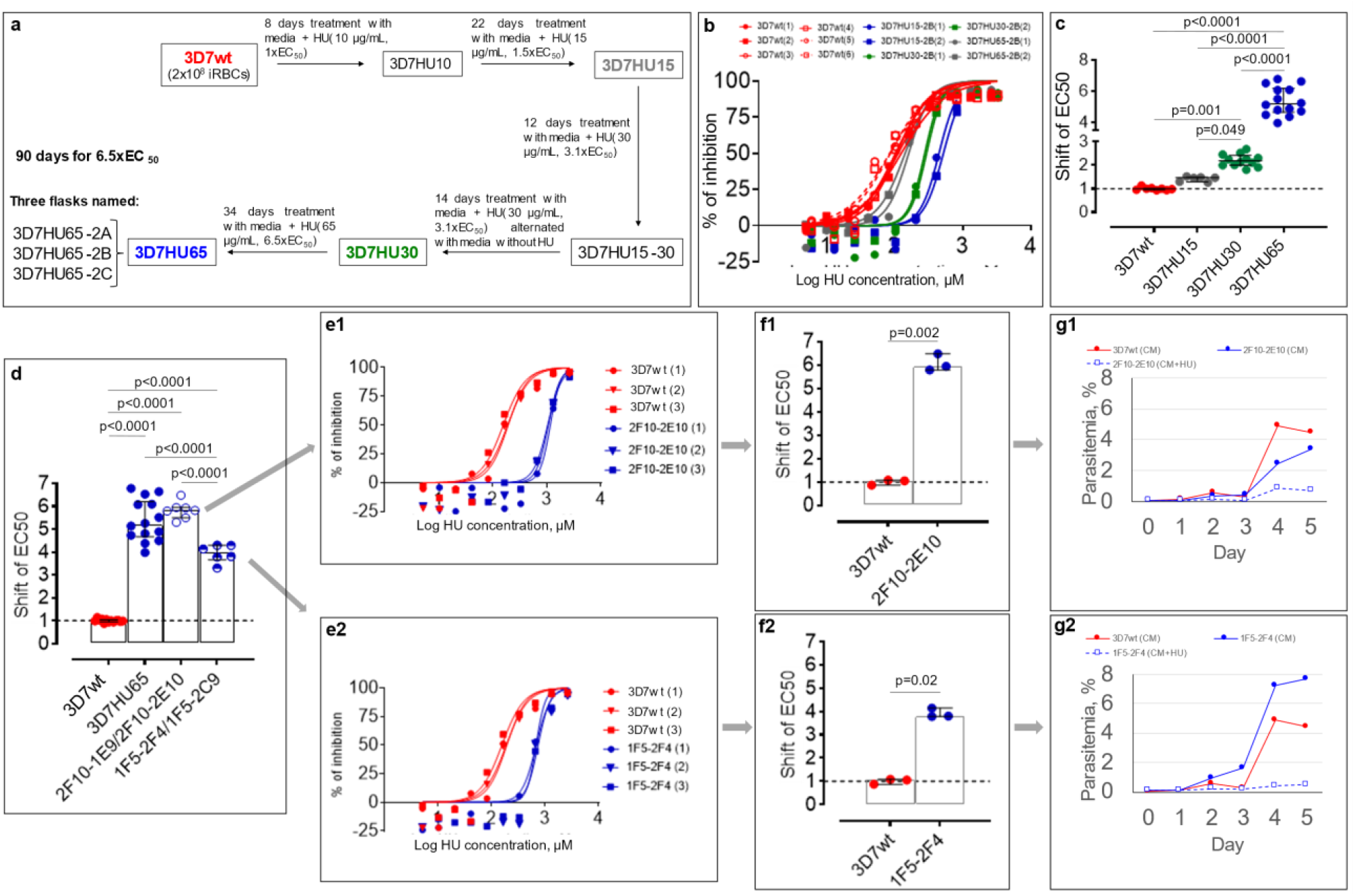
*In vitro* selection of HU-resistant *Pf*3D7 strain. **a** Diagram of selection schemes. Stepwise selections: Three independent parasite cultures were exposed to stepwise increasing concentrations of HU (131.5 to 854.65 µM) as indicated. **b** A representative dose-response curves of HU against parental strain (3D7wt) and bulk-selected HU resistant parasite populations at different time-point of the selection (3D7HU15, 3D7HU30 and 3D7HU65). **c** Shift in EC_50_ of bulk-selected HU exposed parasite populations at different time-point of the selection (3D7HU15, 3D7HU30 and 3D7HU65). **d** Shift in EC_50_ of the bulk-selected 3D7HU65 parasite populations and four clones with higher (2F10-1E9, 2F10-2E10) or lower (1F5-2F4 and 1F5-2C9) EC_50_. **e** Dose-response curves of clones 2F10-2E10 (Panel e1) and 1F5-2F4 (Panel e2) and parental *Pf*3D7 (3D7wt). **f** Shift in EC_50_ of clones 2F10-2E10 (Panel f1) and 1F5-2F4 (Panel f2). **g** Growth response of clones 2F10-2E10 (Panel g1) and 1F5-2F4 (Panel g2) maintained in media without HU (CM) or with HU at 6.5x EC_50_ (CM+HU65). Parental *Pf*3D7 strain (3D7wt) also shown. In panels c, d and f, the horizontal bar represents the median and vertical bar the interquartile range. For the mean comparisons, one-way ANOVA with a Tukey post hoc analysis was used in panels c and d, and Mann-Whitney test in panels f.

Since HU inhibits RNRs^30^ the most likely target in *P. falciparum* is RNR class I which is a complex of homo-dimers of two large (*Pf*R1, chain α) and two small (*Pf*R2, chain β) subunits^31^. The α chain is encoded by *pfr1* gene located in chromosome 14, while the β chain is encoded by two genes *pfr2 and pfr4* located in chromosome 14 and 10, respectively^31 32^. Both large and small subunits are required for activity ^33^. The α chain accommodates the allosteric regulatory sites and the catalytic sites, while the β chain houses the diferric-tyrosyl radical that is essential for catalysis ^32 33^. We therefore examined the effects of HU on *pfr1, pfr2* and *pfr4* genes. We also investigated the multidrug resistance 1 gene (*pfmdr1*) known to be involved in the mechanisms of resistance to a variety of drugs. Analyses were conducted with the trophozoite stage. As shown in Fig. 6a, qPCR analyses suggest no change in expression of *pfr1* in HU-resistant *Pf*3D7 (Fig. 6a). In contrast, the genes coding for the small subunit of RNR enzyme (*pfr2* and *pfr4*) and *pfmdr1* were significantly increased in HU resistant parasites compared to the *Pf*3D7 parent line (Fig. 6a). The level of HU resistance of 3D7HU65 (median, interquartile range value: 5.2, 4.7 – 6.2) was better correlated with the expression of *pfr2* (3.8, 3.0 – 5.3) and *pfr4* (4.7, 3.3 – 7.9) genes compared to the *pfmdr1* gene (2.7, 2.5 – 3.0) (Fig. 6a), suggesting over expression of *pfr2* and *pfr4* genes may contribute to HU resistance of 3D7HU65 parasites.

**Fig. 6.**
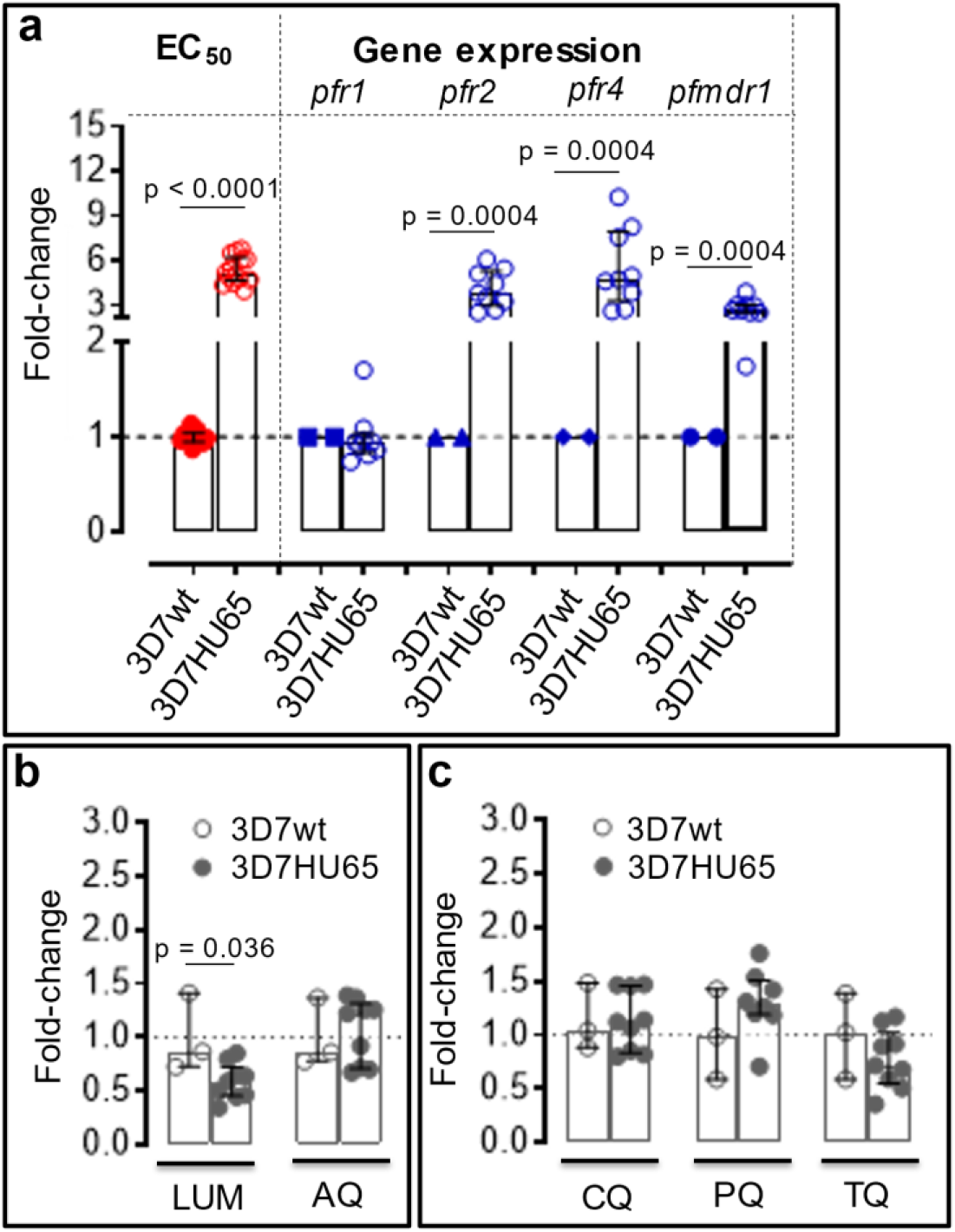
Correlation between HU-resistance gene-expression profile for bulk-selected, HU tolerant 3D7HU65 parasites and efficacy of ACT partner drugs against HU-tolerant strains. **a** Correlation between change in EC_50_ and levels of gene expression of RNR (*pfr1, pfr2* and *pfr4*) and multidrug resistance marker 1 (*pfmdr1*) genes of HU-tolerant 3D7HU65 parasites analyzed by qRT-PCR. Parental *Pf*3D7 line (3D7wt), was used as baseline. **b-c** Anti-plasmodial activity of current antimalarial drugs against 3D7HU65 HU-tolerant parasites. The horizontal bar represents the median and vertical bar the interquartile range. For the mean comparisons, Mann-Whitney test was used for fold-change of EC_50_ in panels a, b and c, and Wilcoxon signed rank test for fold-change of gene expression in panel a. LUM, lumefantrine; AQ, amodiaquine; CQ, chloroquine; PQ, primaquine and TQ, tafenoquine.

Although we were unable to select for *P. falciparum* stably resistant to HU, we utilized 3D7HU65 parasites that were tolerant of the drug to test their killing by five anti-malarial partner drugs used in ACTs, namely LUM, AQ, CQ, PQ and TQ. As shown in Fig. 6b-c, all of the antimalarials effectively killed HU-tolerant parasites (Fig. 6b-c). Notably, LUM was more effective against HU-resistant parasites compared to the parental *Pf*3D7 line (Fig. 6b).

Taken together these data suggest that *P. falciparum* is refractory to developing resistance to HU. Moreover, current antimalarials remain effective against HU-tolerant parasites (and one LUM, the most widely used ACT partner drug) is significantly more active against HU-tolerant parasites) confirming that exposure to HU does not result in resistance to antimalarials but may increase susceptibility to an important ACT partner drug.

### A model for simultaneous treatment of a genetic disorder that co-evolved with a pathogen

Our data strongly support the expanding long-term use of HU in malaria endemic countries to treat SCA and enhance malaria elimination. They also suggest a generalizable model for simultaneous treatment of genetic disorders that co-evolved with human pathogens. As summarized in Fig. 7, if treatment for a genetic disorder like SCA did not antagonize malaria infection (Fig. 7, treatment 1) the patients would remain highly susceptible to severe malaria and death and thereby possibly compromise even short-term gains made with treating SCA. A treatment with modest anti-pathogen activity that bypasses resistance, effects long-term reduction of the pathogenic load presents a strategy of choice (Fig. 7, treatment 2). A treatment that was potent against infection (Fig. 7, treatment 3) would expectedly lead to resistant parasites proliferating in infected patients and preclude long-term treatment (of months to years) needed for the genetic disorder. Interactions with current drugs to the infection would also be important. Antagonist interactions present challenges (Fig. 7 treatment 4) while favorable drug-drug interactions combined with modest anti-infection activity as seen between HU and antimalarials, present the optimal path (Fig. 7, treatments 3&5). This in conjunction with improvement of the host genetic disorder (which in the case of SCA is through stimulation of hematopoiesis and reducing vasculature inflammation) yields a safe long-term treatment that can be deployed in global health.

**Fig. 7.**
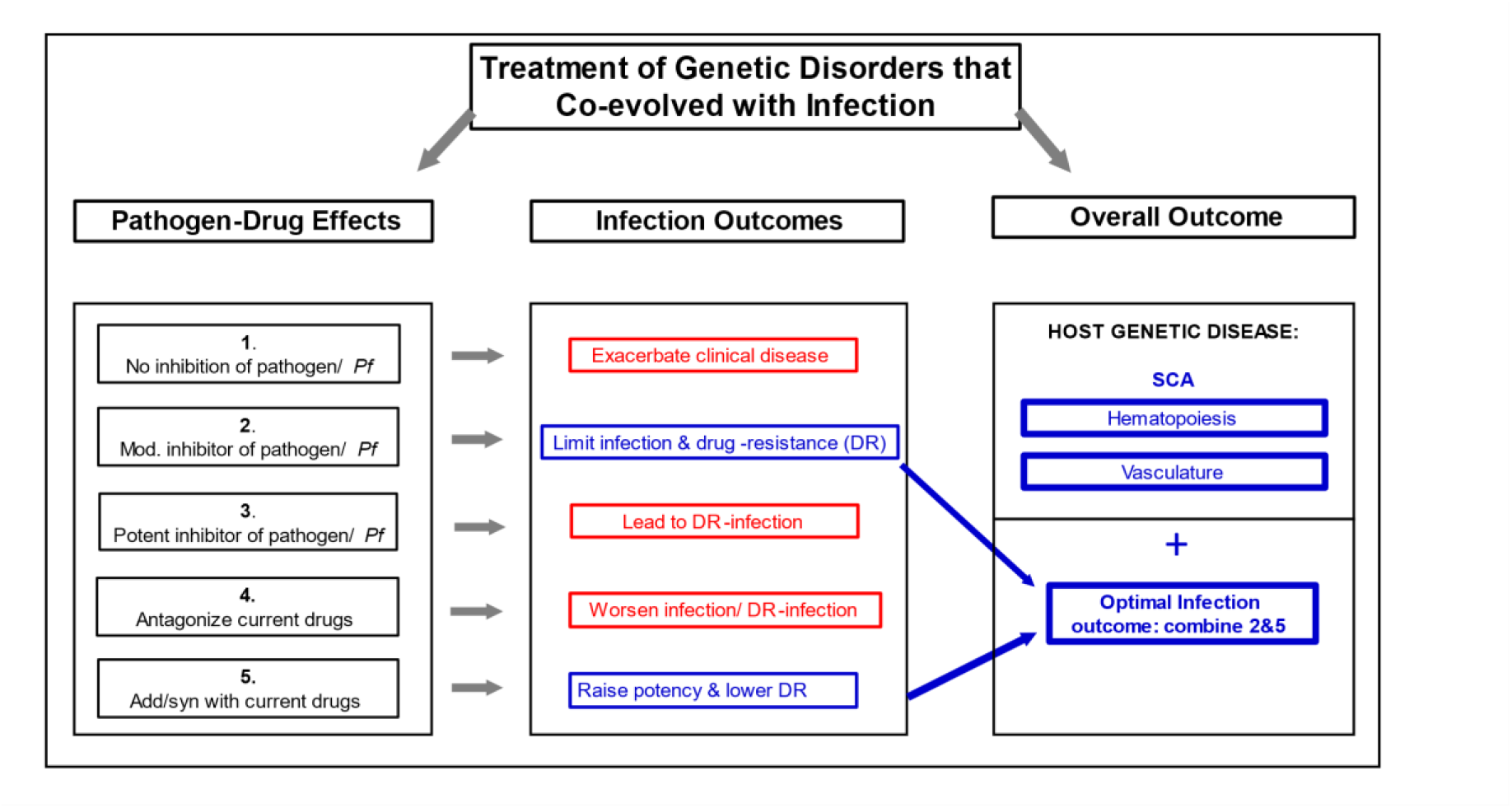
A biological model for treatment of a genetic disorder that co-evolved with infection. In this model infection outcomes are assessed for treatments of the host genetic disorder. Treatment 1 which have no effect on malaria, would leave SCA patients highly susceptible to severe malaria and death. Treatment 2 with modest anti-infection activity that bypasses resistance, presents a strategy of choice. Treatment 3 potent against infection are expected to select resistant parasites given long-term treatment (of months to years) needed to treat the genetic disorder. Treatment 4 with antagonist interactions with antimalarial drugs are undesirable, while favorable drug interactions (Treatment 5) combined with modest anti-infection activity as seen between HU and antimalarials (Treatment 2), present the optimal path. This in conjunction with improvement of the host genetic disorder (which in the case of SCA is through stimulation of hematopoiesis and reducing vasculature inflammation) yields a safe long-term treatment that can be deployed in global health.

## Discussion

Infectious diseases are hypothesized to play a critical role in the selection and maintenance of defective genetic alleles within the human gene pool. Malaria has played a profound role in the development of multiple genetic disorders affecting the red blood cells, but especially for the inherited haemoglobinopathies; the protective effects of sickle trait against severe malaria and death due to malaria are well established. Notably, SCA patients have lower levels of parasite burden than those with normal hemoglobin^16 17^. In this setting, why effective HU treatment of homozygous SCA individuals, did not increase but rather significantly reduced *P. falciparum* malaria in the REACH trial, is poorly understood. Further, the potential interactions of HU with widely used antimalarials, particularly ACTs that have played an important role in malaria elimination, remained unexplored. Our findings suggest that repeated daily exposure to HU alone, at levels achieved in human plasma (even in absence of specific antimalarial treatment) administered over months to years, may be sufficient to kill *P. falciparum* malaria and reduce clinical malaria infections in children with SCA.

Prior studies have reported results on the effects of HU on parasite-induced cerebral malaria in mouse models^34^ and minor killing activity of *P. falciparum in vitro*^35^. Importantly, these studies assessed effects of continuous HU exposure since the pharmacological levels of exposure of HU in patients was not defined. Furthermore, the parasite developmental stage at which HU acts was not defined. We utilized the ‘pulse’ method to better mimic pharmacologic HU exposure in patients with SCA and also discovered stage specific killing effects of HU. The high HU EC_50_ of 127.6 µM, contrasts the EC_50_ of current antimalarials in the low to mid-nanomolar range. This, along with HU’s preferred killing of schizonts and median **∑**FICs achieved in drug-drug interactions, support the hypothesis that HU acts additively with antimalarials rather than changing their intrinsic potency. Importantly HU fails to antagonize a broad range of effective antimalarials and multiple parasite strains, further suggesting that its deployment would not hinder malaria elimination in SCA patients across multiple countries and continents, despite country-specific differences in use of antimalarials.

Prolonged exposure to potent antimalarials (acting at low nanomolar concentrations) has led to emergence of *P. falciparum* resistant to major antimalarials both *in vivo* and *in vitro*. The relatively high EC_50_ for HU of 127.6 µM is the likely reason underlying our observations that *P. falciparum* parasites are refractory to developing stable resistance to HU, even after exposure at concentrations and times that vastly exceed those achieved by the drug in human plasma. We were able to detect HU-tolerant parasites that arise associated with amplification of RNR, mdr and possibly other unidentified genes. However, even if such HU-tolerant parasites were to transiently appear *in vivo*, our findings suggest that they would be rapidly killed by a wide range of antimalarials that are partner drugs of ACTs. Since LUM is significantly more active against HU-tolerant parasites, the tolerance may come at cost to the parasite, a hypothesis that is further strengthened by our data that stable resistance could not be established.

Adjunctive drug-action on the infection and low risk of resistance selection, coupled with treatment of its co-evolved genetic disease, is novel and may be unique for malaria and SCA. But as summarized in Fig. 7, the paradigm may also conceptually be generalized to guide deployment of a broader range of genetic treatments that also improve infection control in global health settings. SCA and β thalassemia are the two common hemoglobinopathies worldwide. There is emerging evidence that HU may be beneficial to the treatment of βthalassemia^36^, which is present in the Mediterranean region, South East Asia and South Asia. Therefore deployment of HU in these regions could provide adjunct therapy of malaria in β thalassemia patients. All major hemoglobinopathies have co-evolved with malaria and thus emerging treatments to these diseases must be evaluated for their effects on plasmodial infection. There is no data yet for other genetic diseases co-evolved with infection, such as cystic fibrosis-cholera, Tay-Sachs-tuberculosis as well as phenylketonuria-mycotic abortions^37^. Nonetheless the model in Figure 7 should be applicable to a broad range of red cell disorders that have co-evolved with malaria, including other common qualitative (HbC and HbE) and quantitative (α thalassemia) hemoglobinopathies.

Finally, the clear additive effects of HU with PQ, suggests the need to evaluate the potential adjunctive effects of HU on *P. vivax*, a second major human malaria parasite species that is on the rise in Africa and worldwide. This would be important for SCA patients but also more broad management of PQ, which is contraindicated in humans with glucose-6-phosphate deficiency, a common enzymopathy affecting 400 million people^38^ that also has co-evolved with malaria.

## Materials and Methods

### *Plasmodium falciparum* culture

Asexual blood stages of three different *P. falciparum* strains 3D7 (*Pf*3D7), NF54 (*Pf*NF54) and Dd2 (*Pf*Dd2) (obtained from BEI resources https://www.beiresources.org/About/MR4.aspx) were cultured in A+ human erythrocytes at 5% hematocrit in RPMI 1640 supplemented with 89 µM hypoxanthine, 25 mM HEPES, 11 µM glucose, 1.74 g/liter sodium bicarbonate, 0.5% (w/v) AlbuMax II (Gibco-BRL) and 24 µg/mL gentamicin in flasks gassed with mixture of 5% O_2_, 5% CO_2_ and 90% N_2_ gas and maintained at 37°C in a humidified atmosphere.

### *In vitro* drug assays

#### Drug preparations

RNR specific inhibitors hydroxyurea (catalog number: H8627-1G), clofarabine (C7495-5MG), gemcitabine hydrochloride (G6423-10MG) and triapine (or 3-AP, SML0568-25MG), and the antimalarial drugs lumefantrine (L5420-5MG), amodiaquin dihydrochloride dihydrate (A2799-5G), chloroquine diphosphate salt (C6628-25G), primaquine diphosphate (P2940000), tafenoquine succinate (SML0396-10MG), quinine hydrochloride dihydrate (Q1125-5G) and dihydroartemisinin (D7439-50MG) were all purchased from Sigma-Aldrich (USA). The stock of RNR inhibitors, lumefantrine, amodiaquine, primaquine, tafenoquine and dihydroartemisinin were prepared in dimethyl Sulfoxide (DMSO), chloroquine in milli Q water and quinine in 70% ethanol. The stocks were kept in -20°C for no more than 6 months.

#### 72 h *in vitro* antimalarial drug assay

*In vitro* effective concentration values were determined by incubating parasites at 0.5% starting parasitemia and 1.5% hematocrit with a range of drug concentrations (across a range of 2-fold serial dilutions) at 37°C for 72 h in flat-bottom 96-well plates in a 100 μl culture volume. Four wells without compound and two wells with uninfected RBCs were used as positive and negative controls, respectively. Outer wells were filled only with distilled water to reduce evaporation in the plate. For stage specificity assays, the starting parasitemia was 0.2% for schizont stage. Parasite growth in each well was assessed after 72 h by flow cytometry^39^. Briefly, 25 μl samples were harvested after 72 hours and stained for 30 min at 37°C in a humidified atmosphere with 25 μl of 1× SYBR Green I (S7563, Thermo Fisher Scientific) plus 300 nM MitoTracker Deep Red (M22426, Thermo Fisher Scientific, USA) diluted in culture medium. Samples were washed in RPMI 1640 and centrifuged for 4 min, 2000 rpm at room temperature (RT). Excess stain was removed, and half of the samples were resuspended in 300 μl of RPMI and analyzed with BD LSRFortessa X-20 flow cytometer. EC_50_, EC_90_ and EC_99_ values were determined by nonlinear regression analysis using GraphPad Prism 9.2.0.

#### Checkerboard method

Ring *Pf*3D7 (age < 9 hours) was exposed to a fixed concentration of hydroxyurea (20.6 to 329 µM) plus varied concentrations of primaquine ranging from 0 to 10 µM in a classical 72 h *in vitro* antimalarial drug assay as described above. The assays were conducted in flat-bottom 96-well plates in a 100 μl culture volume, 1.5% hematocrit, and 0.5% starting parasitemia. The parasitemia was determined by flow cytometry using a BD LSRFortessa X-20 flow cytometer with parasites stained with SYBR green I and MitoTracker Deep Red as described above.

#### Pulsed method

Highly synchronized parasites were pulsed 3 or 6 hours at 37°C with 329 µM hydroxyurea or 0.1% DMSO in 500 µl of complete media (0.5% staring parasitemia, 1,5% hematocrit), then transferred into 15 ml polystyrene tube, centrifuged for 5 min, 2000 rpm at RT, washed twice in 5 ml of warm RPMI-1640 and incubated in complete media with or without drug, or with 0.1% DMSO for 69 or 66 hours. At the end of incubation time, the parasitemia was quantified by flow cytometry as described above. The starting parasitemia was 0.2% for schizonts.

#### Fixed-ratio method

Drug interaction studies were performed using a modification of the fixed ratios method^27^. Dose-response assays were first carried out to obtain the EC_50_ values of the individual test compounds. Subsequently, hydroxyurea and the antimalarial compounds were diluted in complete culture medium to initial concentrations of 10 times the predetermined EC_50_ and the solutions combined in ratios of 9:1, 7:3, 5:5, 3:7 and 1:9. Serial two-fold dilutions (10 concentrations) were then prepared in these fixed ratios. 50 µl/well diluted drug mixture were added to 50 µl/well of infected RBCs (0.5% parasitemia) and dispensed into 96-well plate (1.5% final hematocrit), then maintained in a humidified atmosphere in incubator at 37°C with 5% CO_2_ during 72 h. Four drug-free control well and 2 wells with uninfected RBCs were also included. Parasite growth in each well was assessed after 72 h by flow cytometry as described above. EC_50_, EC_90_ and EC_99_ values of each compound alone and in combination were determined by nonlinear regression analysis using GraphPad Prism 9.2. For data interpretation, the EC_50_ or EC_90_ or EC_99_ of the drugs in combination (hydroxyurea and antimalarial drug) were expressed as fractions of the EC_50_ or EC_90_ or EC_99_ of the individual drugs. These fractions were called fractional inhibitory concentrations (FIC) for hydroxyurea or FIC(HU) and for antimalarial drug or FIC(antimalarial drug). Isobolograms were constructed by plotting the FIC(HU) against the FIC(antimalarial drug) for each of the five ratios, with concave curves indicating synergy, straight lines indicating addition and convex curves indicating antagonism. To obtain numeric values for the interactions, results were expressed as the sum FICs (∑FICs) of the FIC(HU) and FIC(antimalarial drug). The interpretation of these numeric values was done as previously described^29^. ∑FIC < 0.5 indicates substantial synergism, ∑FIC <1 indicates moderate synergism, ∑FIC ≥1 and <2 indicates additive interaction, ∑FIC ≥2 and <4 indicates slight antagonism and ∑FIC > 4 indicates marked antagonism. Median ∑FICs were used to classify the overall nature of the interaction.

#### *In vivo* drug assay

Animal protocols were reviewed and approved by the institutional animal care and use committee (IACUC) of University of Notre Dame (Approved Protocol # 21-03-6520). The effect of hydroxyurea on anti-plasmodial activity of primaquine or tafenoquine was tested in a mouse model of *P. berghei* infection using the 4-day suppressive test according to the procedure detailed by Fidock *et al*.^40^. Briefly, C57BL/6 mice were first injected intraperitoneally (i.p.) with thawed stabilates of blood-stage *P. berghei* ANKA (obtained from bei https://www.beiresources.org/About/MR4.aspx). Subsequently, live parasites isolated from these mice were used to infect 6 to 8 weeks old male or female C57BL/6 mice, weighing 16.2 – 28.4 g by inoculating i.p. 2 × 10^7^ *P. berghei* ANKA infected red blood cells. Mice were randomly divided into four or five groups of 3 to 5 mice each. Four single doses of HU (50 mg/kg) and/or primaquine (0.1 or 1 mg/kg) solubilized in phosphate buffer saline (PBS, vehicle) or just vehicle was administered i.p. starting 2-4 hours after infection and every 24 hours for 3 days. The stock solutions of the drugs were prepared in the vehicle to the desired final concentration so that each animal received between 150 to 200 μl at time of administration of each drug. Twenty-four hours after the last treatment (i.e. 96 hours’ post-infection), animals’ weight was compared to original body weight, and the parasitemia analyzed by Giemsa-stained tail blood smears and flow cytometry as previously described^41^. Briefly, 2.5 μl of tail blood were stained for 20 min with a slightly agitation at RT in 150 μl of filtered PBS-BSA (0.5%) containing 0.25× SYBR Green I, 8.5 μg/ml dihydroethidium (D11347, Thermo Fisher Scientific) and 1.33 μg/ml CD45 rat anti-mouse, PE-Cy7 (clone 30-F11, 25-0451-82, Thermo Fisher Scientific). After incubation, 850 μl of cool PBS 1x were added and centrifuge 2000 rpm, 3 min at RT, then the supernatant was discarded, and the pellet suspended in PBS-BSA (0.5%) and analyzed with BD LSRFortessa X-20 flow cytometer.

For HU and TQ combination, the study design was similar to that used for HU and PQ with some modifications regarding the posology of TQ. For the first experiment, a single treatment with tafenoquine (0.5 mg/kg) was given 2-4 hours after infection but in two experiments, four treatments with TQ (0.5 mg/kg) were given as indicated above.

#### *In vitro* selection of *P. falciparum* resistant to HU

We adapted well established methods with two strains *Pf*3D7 and *Pf*Dd2^42^. Briefly we began with single-step selections undertake in triplicate flasks each containing 2 × 10^9^ parasites exposed to HU at 3× or 6× the EC_50_ (394.5 or 789 µM for *Pf*3D7; 678 or 1315 µM for *Pf*Dd2). Viable parasites failed to emerge with either strain after 60 days and the selections were therefore discontinued.

We next used ramping selections in triplicate flasks of 2 × 10^8^ parasites that were exposed to gradually increasing concentrations of HU. We used 1× to 6× the EC_50_ (131.5 to 855 µM) over 3 months for *Pf*3D7 and 263 to 1315 µM HU for 8 months for *Pf*Dd2). Surviving parasites that emerged in ‘bulk’ cultures were cloned by two rounds of limiting dilution in drug-free media in 96-well plates (containing on average 0.15 parasite per well). These plates were screened for viable parasites after 14 to 21 days by staining 20 μl/well of samples with SYBR Green I and Mito Tracker Deep Red followed by flow cytometry (BD LSRFortessa X-20 flow cytometer) as described above. Isolated clones were expanded and 4 were selected for phenotypic and biological characterization.

#### Quantitation of *P. falciparum* RNR and *mdr1* gene expression and copy number by real-time PCR

Primers from *pfr1* (GenBank accession no. U01323.1), *pfr2* (GenBank accession no. U01322.1) and *pfr4* (GenBank accession no. AY669809.1) genes were designed with Primer Express software 3.0 (Applied Biosystems). The primers and probes from *pfmdr1* and *β-tubulin* genes (Supplementary Table 1) were as previously reported.

Parasite genomic DNA and total RNA template were extracted from parasite culture using Quick-gDNA™ Blood MiniPrep (D3072, Zymo Research) and QIAamp RNA Blood Mini Kit (52304, QIAGEN), respectively according to the manufacturer’s instructions. DNA and RNA extracts were quantified using a NanoDrop ND-1000 (NanoDrop Technologies, Wilmington, DE, USA). **SYBR Green I RT-PCR**. Gene expression was analyzed by reverse transcription-polymerase chain reaction (RT-PCR). The assay was performed using the Applied Biosystems StepOnePlus Real-time PCR system with a 20 µL reaction volume containing 1ng of total RNA and the SYBR® Green PCR Master Mix (4309155, Thermo Fisher Scientific) in MicroAmp® Fast Optical 96-Well Reaction Plate (4346906, Thermo Fisher Scientific) using the default thermocycler program for all genes: 30 minutes of reverse transcription at 48°C, then 10 minutes of pre-incubation at 95°C followed by 40 cycles for 15 seconds at 95°C and one minute at 60°C. Melting curve analysis was always performed at the end of each RT-PCR assay to control for specificity of the amplification. **TaqMan PCR**. *pfmdr1* copy number was assessed by TaqMan real-time PCR (Applied Biosystems StepOnePlus Real-time PCR system) as previously described^43^. Primers and fluorescence-labeled probes were used to amplify *pfmdr1* and β-*tubulin* genes. PCR conditions and thermal cycling conditions were used as described. The *Pf*3D7 and *Pf*Dd2 containing one and four *pfmdr1* copies, respectively, was used as the reference DNA sample.

For both SYBR Green I RT-PCR and TaqMan PCR, parasite strains were analyzed in a minimum of 2 independent experiments, with duplicate wells for each experiment. Serial dilutions of *Pf*3D7 total RNA and DNA (ranging from 8 to 5000 pg/µl) with nuclease-free water (AM9935, Life Technologies) were used to generate a relative standard curve for *pfr1, pfr2, pfr4, pfmdr1* and *pf*-*β-tubulin* in order to determine the amplification efficiencies which must be close to 100% and approximately equal between the target and reference genes^44^. Gene expression and *pfmdr1* copy number was determined by relative quantification between the target (*pfr1, pfr2, pfr4* and *pfmdr1*) and the house-keeping (*pf*-*β-tubulin*) genes (for gene expression analysis), or between *pfmdr1* and *pf*-*β-tubulin* genes (for *pfmdr1* copy number analysis) based on the ΔΔCt method as previously described^44^. In brief, the target gene (*pfr1, pfr2, pfr4* and *pfmdr1*) and reference (*pf*-*β-tubulin*) genes were amplified with the same efficiency within an appropriate range of RNA or DNA concentrations. The relative expression or copy number of the target genes was calculated using the 2^−ΔΔCt^ method where ΔΔCt = (Ct, target gene – Ct, *Pf*-β-*tubulin*)χ – (Ct, target gene – Ct, *Pf*-β-*tubulin*)y, with χ = HU resistant parasite or *Pf*Dd2 and y = *Pf*3D7. The result is expressed as n-fold change of target gene expression or copy number relative to that of *Pf*3D7.

### Statistical analysis

The Mann-Whitney U test, Wilcoxon signed-rank test or one-way analysis of variance (ANOVA) with a Tukey post hoc analysis was used for mean comparisons using GraphPad Prism 6 software.

## Data Availability

Accession numbers for primers are indicated as follows: *pfr1* (GenBank accession no. U01323.1), *pfr2* (GenBank accession no. U01322.1) and *pfr4* (GenBank accession no. AY669809.1) The authors declare that all data supporting the findings of this study are available withing the paper and its Supplementary information. Requests for resources and reagents should be directed to KH.

## Acknowledgements

This work was supported by the National Institutes of Health USA, R01 HL 1330330 (KH). REW receives relevant grant funding from the National Institutes of Health (U01 HL133883) and Doris Duke Charitable Foundation (2019163).

## Author Contributions

IS. Conceptualization, study design, execution of all experimental studies, curation and analyses of all data, including statistical analyses, visualization of results, drafting and editing of manuscript. RW. Conceptualization and study design, data curation and analyses, visualization of results, drafting and editing of manuscript.

NM. Conceptualization and study design, data curation and analyses, visualization of results, drafting and editing of manuscript.

KH. Principal Investigator, overall conceptualization and project supervision and design; curation and analyses of all data, visualization of results; drafting and editing of manuscript.

## Inclusion and Ethics

The authors declare no competing interests.

