## Supplemental Information for "Simultaneous adjunctive treatment of malaria and its co-evolved genetic disorder sickle cell anaemia"

### Supplementary Table and Figure

### Supplementary Table 1 Primer and probe sequences

|  | Primer | **Sequence** (5'----3') |
|---|---|---|
| SYBR Green I RT-PCR | *pfr1*-F | ATA TGT AGA TCA AGG TGG AGG AAA AC |
| | *pfr1*-R | CTC TCT TCA TAA AAA GAT CAG GAA CC |
| | *pfr2*-F | TTA GGA TGC TCT AAA ATT TTC CAT TC |
| | *pfr2*-R | AAA ATT CCG TAT TCA GAC AAA AGA CT |
| | *pfr4*-F | GAT ACC TGA AGG AAG AGC TTT TTA TG |
| | *pfr4*-R | ATT CGA ATA TTT CAG TTA GCC ATT TC |
| SYBR Green I RT-PCR and Taqman PCR | *pfmdr1*–1F* | TGC ATC TAT AAA ACG ATC AGA CAA A |
| | *pfmdr1*–1R* | TCG TGT GTT CCA TGT GAC TGT |
| | *pfmdr1*-probe* | FAM-TTT AAT AAC CCT GAT CGA AAT GGA ACC TTT G-TAMRA |
| | *β-tubulin*-1F* | TGA TGT GCG CAA GTG ATC C |
| | *β-tubulin*-1R* | TCC TTT GTG GAC ATT CTT CCT C |
| | *β-tubulin*-probe* | VIC-TAG CAC ATG CCG TTA AAT ATC TTC CAT GTC T-TAMRA |

RT-PCR, Reverse Transcription-Polymerase Chain Reaction; *Designed by Price *et al*.[38]

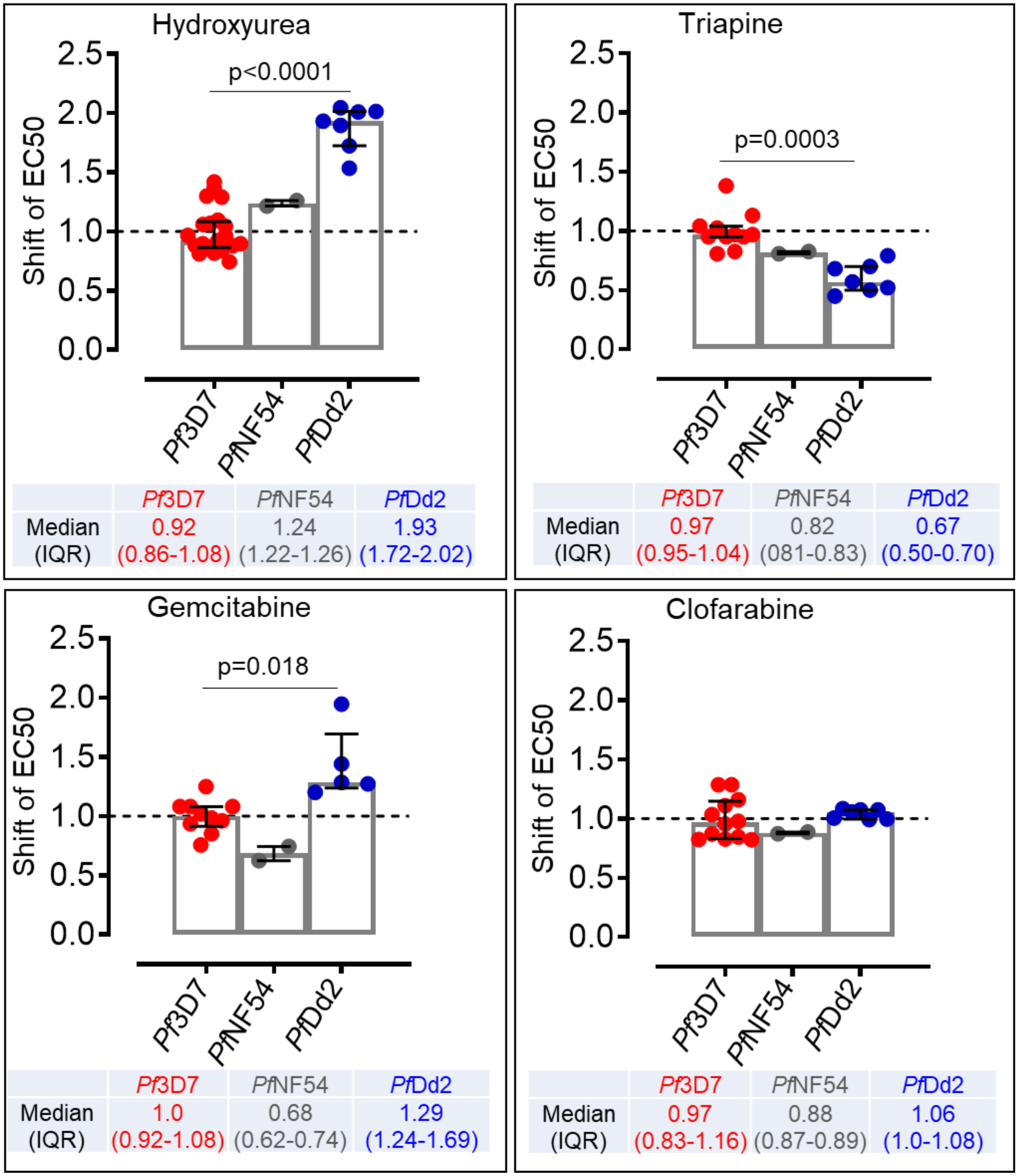


**Supplementary Fig. 1 Fold-change of $EC_{50}$ of *Pf*NF54 and *Pf*Dd2 relative to that of *Pf*3D7 after exposition to HU and other RNR inhibitors *in vitro*.** The horizontal bar represents the median and vertical bar the interquartile range. Kruskal-Wallis test with

Dunn's multiple comparison test was used for the mean comparisons. IQR, interquartile range.

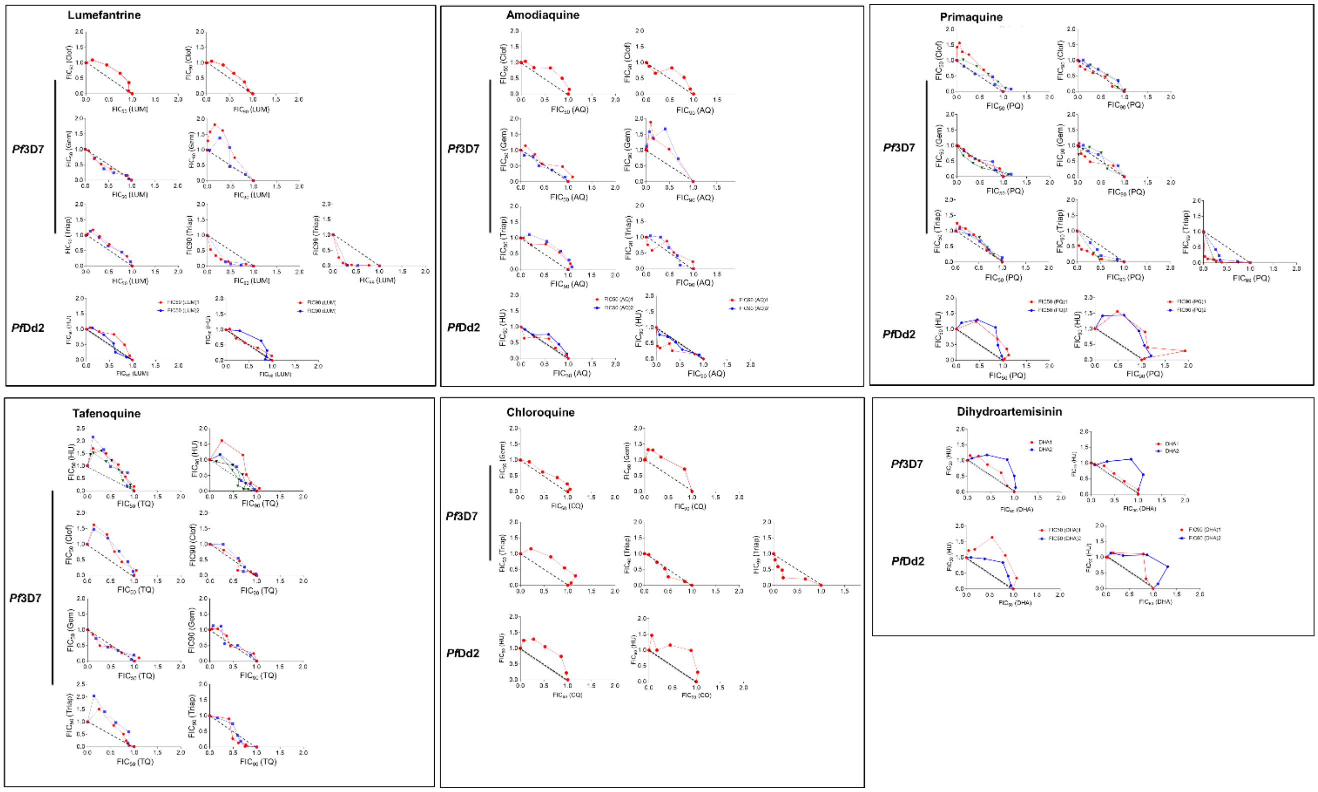


**Supplementary Fig. 2 Isobologram analyses of interactions between HU or other RNR inhibitors and indicated antimalarial drugs against *Pf*3D7 and *Pf*Dd2 strains**. Fractional inhibitory concentrations (FIC) of HU, or other RNR inhibitors *versus* FIC of LUM, AQ, CQ, PQ, TQ and DHA constructed from $EC_{50}$ or $EC_{90}$ or $EC_{99}$ values. For each drug combination, the FIC were calculated by dividing the measured ‘‘apparent’’ EC values for individual drugs in the different combinations of HU or other RNR inhibitors and antimalarial drugs by the EC values obtained when the drugs were used alone. 1 to 5 independent experiments**.**

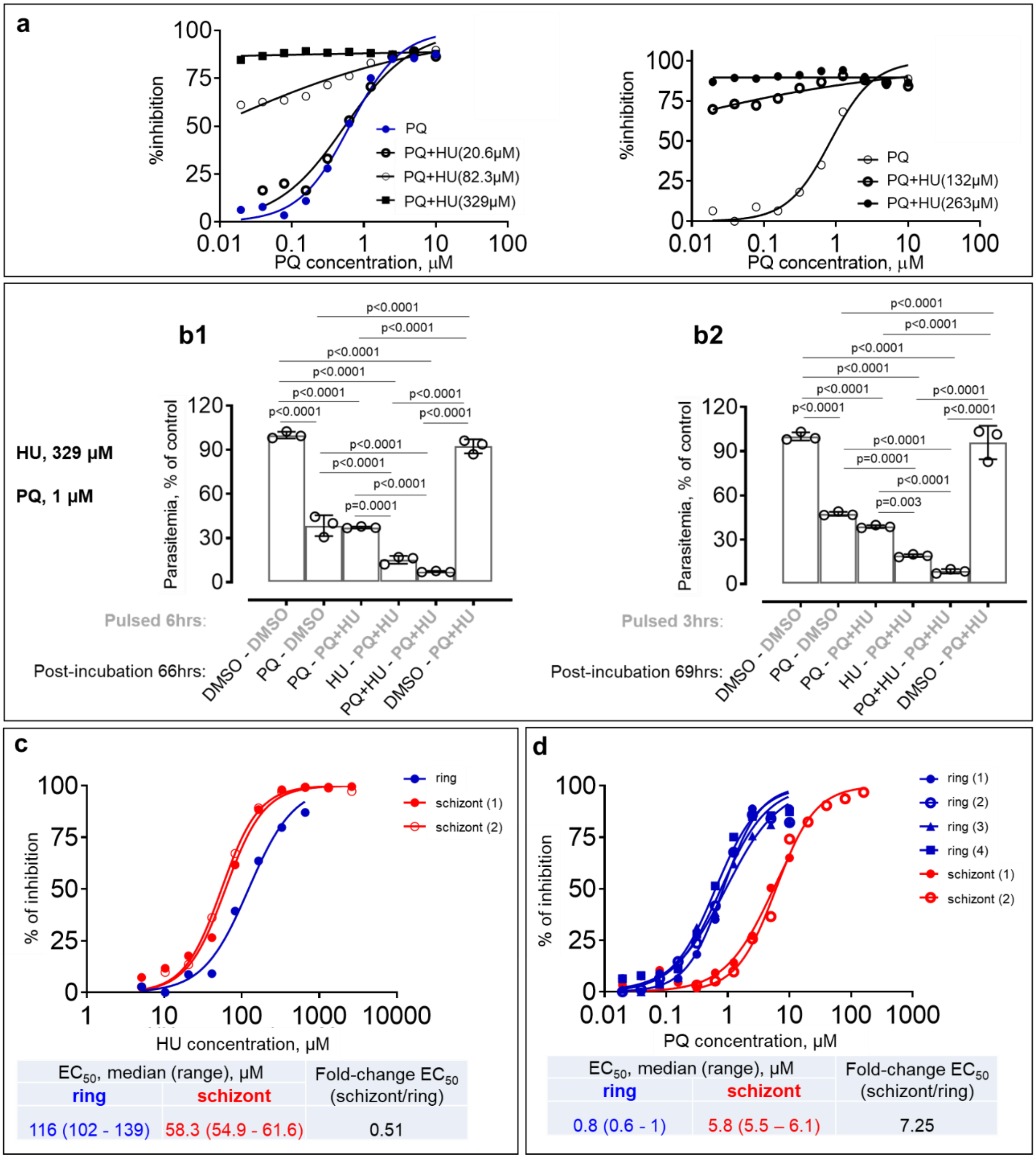


**Supplementary Fig. 3 Potentialization of anti-plasmodial activity of PQ by HU *in vitro*. a** Dose-response curves of PQ + HU against *Pf*3D7 asexual blood parasites starting at early ring stage using checkerboard method. Parasites were exposed to a fixed concentration of HU (20.6 - 329 µM) plus varied concentrations of PQ (0 - 10 µM) for 72 hours. **b** Inhibition rate of PQ and/or HU against ring stage of parasite. *Pf*3D7 ring (age: < 8 hours) were pulsed 6 (Panel b1) or 3 (Panel b2) h with 329 µM HU plus 1 µM PQ, or with 0.1% DMSO, then extensively washed and incubated in media containing 329 µM HU and/or 1 µM PQ, or with 0.1% DMSO for 66 (Panel b1) or 69 (Panel b2) h, respectively. **c-d** Anti-plasmodial activity of HU and PQ against *Pf*3D7 ring and schizont. Dose-response curves of HU (Panel c) or PQ (Panel d) against ring and schizont stages. In panels b, DMSO-treated parasites were used as control. Horizontal bar represents median and vertical bar the interquartile range in panels b. For the mean comparisons in panels b, one-way ANOVA with a Tukey post hoc analysis was used.

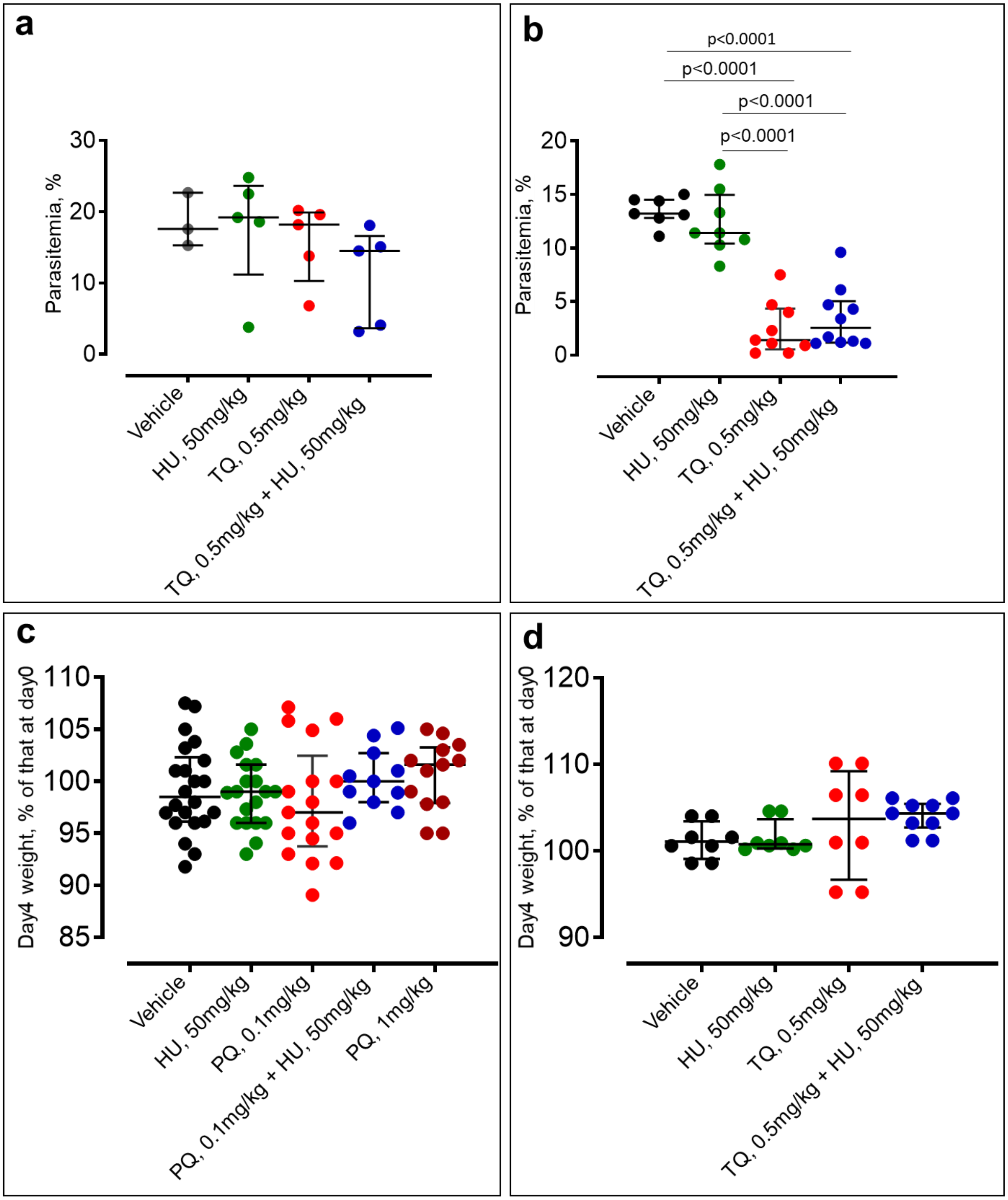


**Supplementary Fig. 4 Anti-plasmodial activity and toxicity of HU combined with TQ or PQ in *P. berghei* Anka-infected mice. a-b** Parasitemia after 4 days of treatment with HU and/or TQ. A single treatment with TQ (0.5 mg/kg) was given 2-4 hours after infection

(Panel a) or four treatments with TQ (0.5 mg/kg) were given starting 2-4 hours after infection and every 24 hours for 3 days (Panel b). **c-d** Weight of *P. berghei* Anka-infected mice after 4 days of treatment with HU alone or in combination PQ (Panel c) or TQ (Panel d). The horizontal and vertical bars represent median and interquartile range. For the mean comparisons in panels b, one-way ANOVA with a Tukey post hoc analysis was used.

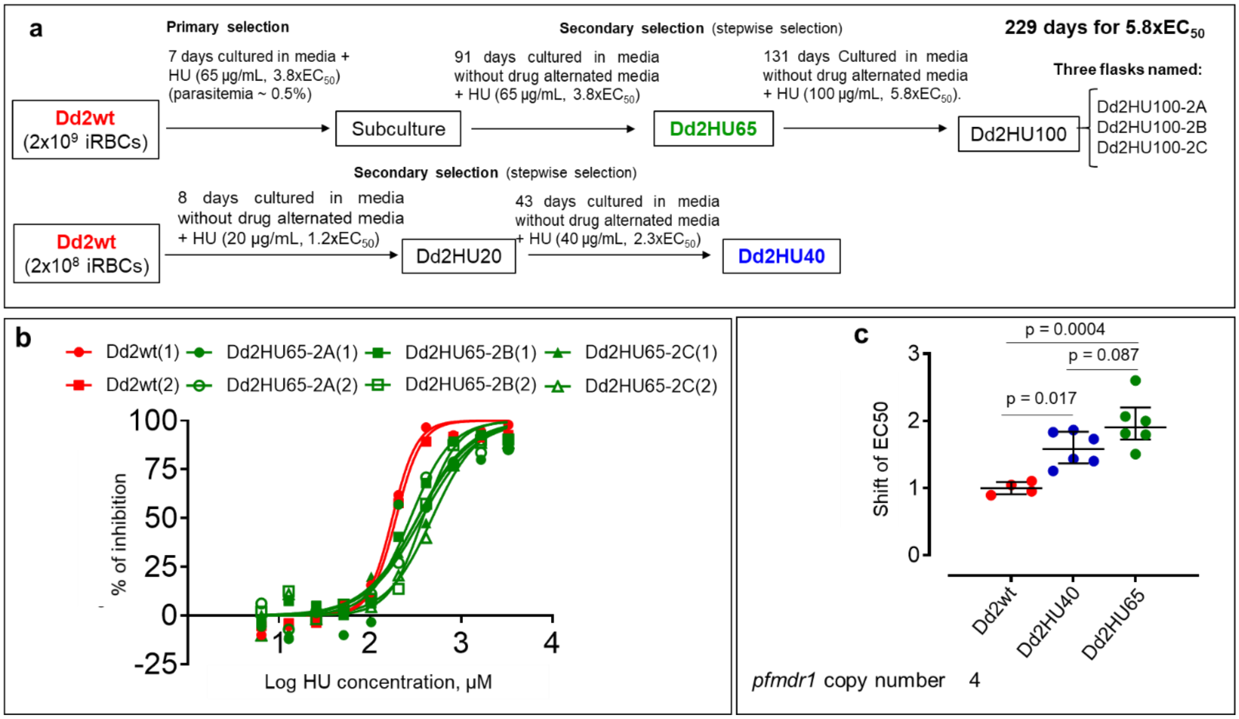


**Supplementary Fig. 5** ***In vitro*** **selection of HU-resistant** ***Pf*****Dd2 strain. a** Diagrams of *in vitro* selection of *Pf*Dd2 resistant to HU. **b** Dose-response curves of HU against parental strain (Dd2wt) and bulk-selected HU-tolerant parasite populations at different time-points of drug pressure (Dd2HU40 and Dd2HU65). **c** Shift in $EC_{50}$ of bulk-selected HU-tolerant parasite populations at different time-point of the selection (Dd2HU40 and Dd2HU65).
